# Characterisation of anhydro-sialic acid transporters from mucosa-associated bacteria

**DOI:** 10.1101/2024.01.17.576040

**Authors:** Yunhan Wu, Andrew Bell, Gavin H. Thomas, David N. Bolam, Frank Sargent, Nathalie Juge, Tracy Palmer, Emmanuele Severi

## Abstract

Sialic acid (Sia) transporters are critical to the capacity of host-associated bacteria to utilise sialic acid for growth and/or cell-surface modification. While N-acetyl-neuraminic acid (Neu5Ac)-specific transporters have been studied extensively, little is known on transporters dedicated to anhydro-sialic acid forms such as 2,7-anhydro-Neu5Ac (2,7-AN) or 2,3-dehydro-2-deoxy-Neu5Ac (Neu5Ac2en). Here, we used a Sia-transport-null strain of *Escherichia coli* to investigate the function of members of anhydro-Sia transporter families previously identified by computational studies. First, we showed that the transporter NanG, from the Glycoside-Pentoside-Hexuronide:cation symporter family, is a specific 2,7-AN transporter, and identified by mutagenesis a crucial functional residue within the putative substrate-binding site. We then demonstrated that NanX transporters, of the Major Facilitator Superfamily, also only transport 2,7-AN and not Neu5Ac2en nor Neu5Ac. Finally, we provided evidence that SiaX transporters, of the Sodium-Solute Symporter superfamily, are promiscuous Neu5Ac/Neu5Ac2en transporters able to acquire either substrate equally well. The characterisation of anhydro-Sia transporters expands our current understanding of prokaryotic Sia metabolism within host-associated microbial communities.

## INTRODUCTION

Sialic acid (Sia) covers a family of over 50 chemically and structurally related 9-carbon sugar acids ubiquitous across vertebrates [1]. Sia residues cap the glycan chain of glycoproteins and glycolipids found on the cell surface of mucosal tissues (such as those lining the respiratory, gastrointestinal, and urogenital tracts), where they mediate key biological interactions based on glycan-protein recognition, regulating numerous processes including immune system functions and neural development [1, 2]. Nacetyl-neuraminic acid (Neu5Ac; Fig. 1) is the most abundant Sia in nature and the only form synthesised *de novo* by humans [3]. Bacteria living in mucosal surfaces can access terminal Sia for their own benefit [4, 5] by incorporating Sia into own cell surface structures (such as capsules and lipopolysaccharides) [5, 6] to evade the immune system by “molecular mimicry” [5] and/or through consumption of Sia for growth to enhance colonisation of established niches [7, 8]. The release of Sia from host surfaces by sialidases and its subsequent uptake into the cell by Sia transporters are essential steps in these processes [4]. Most bacterial sialidases belong to glycoside hydrolase family 33 (GH33) of the CAZy database (www.cazy.org [9]), where they liberate monomeric Neu5Ac from a variety of glycans [4, 10]. In addition intramolecular *trans* (IT)-sialidases from GH33 have been shown to release anhydro-forms of Neu5Ac (Fig. 1) upon cleavage through an intramolecular *trans*glycosylation reaction, leading to 2,7-anhydro-Neu5Ac (2,7-AN) [11, 12] made by e.g., *Streptococcus pneumoniae* D39 NanB [11, 13] and *Ruminococcus gnavus* ATCC 29149 NanH [12], or 2-deoxy-2,3-dehydro-Neu5Ac (Neu5Ac2en) made by e.g., *S. pneumoniae* TIGR4 NanC [14]. The importance of Neu5Ac and 2,7-AN metabolism in niche colonisation has been established in experimental models [4, 7, 15], while Neu5Ac2en metabolism has only been studied *in vitro* using bacterial cultures [16] and/or purified enzymes [14, 16].

**Figure 1.**
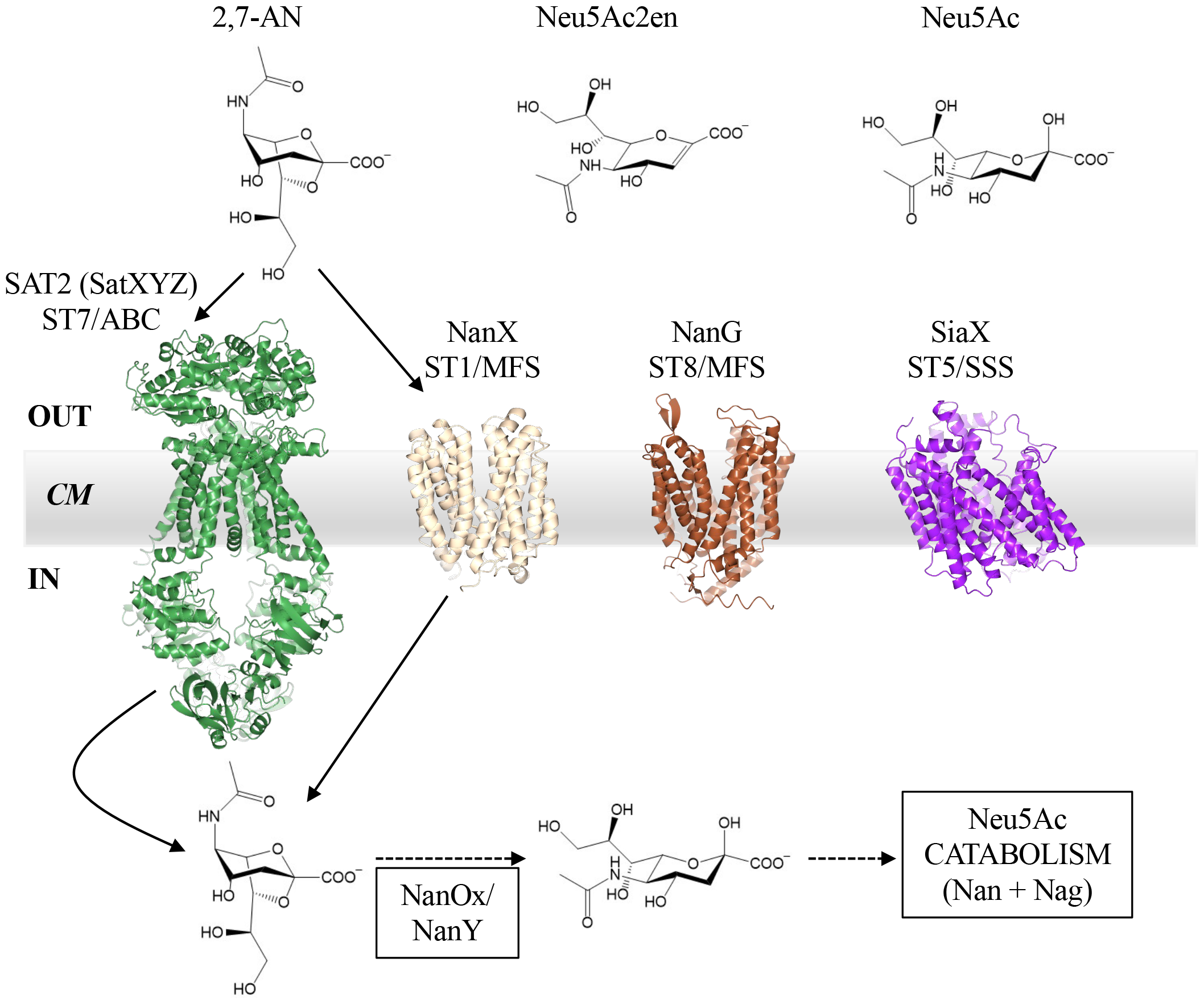
Anhydro-Sia transporters from mucosa-associated bacteria. The four phylogenetically distinct families (ST) of anhydro-Sia transporters are here represented by SAT2 (aka SatXYZ; ST7/ABC) from *R. gnavus* ATCC 29149, NanX (ST1/MFS) from *E. coli* BW25113, and NanG (ST8/MFS) and SiaX (ST5/SSS), both from *S. lutrae* DSM 10244. Sia ABC transporters are multidomain complexes comprising an extracytoplasmic solute-binding protein, a dimeric transmembrane domain, and a dimeric cytoplasmic ATPase domain which energises transport by ATP hydrolysis [43]. In contrast, Sia MFS and SSS systems are secondary (i.e., ion gradient-dependent) transporters comprising a single protein domain, made of structural repeats in either case, but with different folds and internal symmetries between repeats [44, 45]. While both from the MFS superfamily, ST1 and ST8 transporters belong to distinct families [17] which function by different mechanisms [44]. As there is no experimental 3D structure of any anhydro-Sia transporter at present, we show here AlphaFold2 [32] predictions to emphasise the major structural differences among these four groups (the AlphaFold2 prediction of the SatXYZ transporter includes the multitask ATPase, MsmK [46], to complete the functional complex). Anhydro-Sia released by IT-sialidases may be either 2,7-anhydro-Neu5Ac (2,7-AN) or 2-deoxy-2,3-dehydro-Neu5Ac (Neu5Ac2en); Neu5Ac is shown as the β-anomer (ca 90% at equilibrium). Experimentally confirmed uptake pathways are limited to those using *Rg*SatXYZ and *Ec*NanX (arrows), both for 2,7-AN acquisition. NanOx (known as NanY in *E. coli*) is the essential cytoplasmic oxidoreductase converting anhydro-Sia into Neu5Ac [7, 21], which is then catabolised by the conserved Nan and Nag enzymes [4]. CM: cytoplasmic membrane.

To acquire Sia, bacteria employ specific transporters located in the cytoplasmic membrane [17, 18] . Numerous types of Sia transporters have been identified, reflecting the ecological importance of this trait in Sia-rich niches [17]. Sia transporters belong to four superfamilies of prokaryotic transporters with major differences in fold, subunit composition, and mode of energisation [18]: the MFS (major facilitator superfamily), SSS (sodium-solute symporter), ABC (ATP-binding cassette), and TRAP (tripartite ATP-independent periplasmic transporter) superfamilies. Phylogenetic differences within these groups split Sia transporters into eight independently evolved families (“ST1-8”) which differ by finer structural-functional properties [17]. A key difference among Sia transporters is their substrate specificity for Neu5Ac and/or 2,7-AN/Neu5Ac2en [17]. Decades of research on Neu5Ac transporters have identified six experimentally confirmed ST families (ST1-6) [17] and provided detailed genetic and mechanistic studies [4, 17] alongside some experimental 3D structures [4, 19, 20]. In contrast, much less is known on the four types of anhydro-Sia transporters (Fig. 1) identified recently based on computational analyses [17, 21], except for the ABC transporter SAT2 from *R. gnavus* ATCC 29149 (ST7; renamed SatXYZ in [17]), which has been shown to be specific for 2,7-AN [7]. As for NanX (ST1/MFS), found in *Escherichia coli* K12 and other Gram-negative bacteria, the orthologue of *E. coli* BW25113 has been shown to transport anhydro-Sia, but not Neu5Ac, using genetics and growth experiments [16, 21]. However, for NanG (ST8/MFS) and SiaX (ST5/SSS), found in *Firmicutes* (*Bacillota*), the evidence remains solely computational, and is based on the conserved genetic link – characteristic of anhydro-Sia transporters [7, 21], between the transporter and the cytoplasmic oxidoreductase NanOx/NanY [17], which is the essential anhydro-Sia catabolic enzyme converting transported anhydro-Sia into Neu5Ac ([7]; Fig. 1).

Here, we used an improved functional expression system based on a fully Sia-transport-null mutant *E. coli* strain to provide experimental evidence for the substrate preference of the previously uncharacterised NanG and SiaX anhydro-Sia transporters.

## RESULTS

### Functional expression in *E. coli* of NanG, SiaX, and NanX anhydro-Sia transporters

*E. coli* K12 strains possess two Sia transporter genes, *nanT*, coding for a well-known Neu5Ac transporter [22–24], and *nanX*, coding for an anhydro-Sia transporter [16, 21]. Here, we generated an *E. coli* strain derived from SEVY3 [19] named TRXC2 (Table 1) which lacks both native transporter genes and is entirely incapable of Sia transport (Fig. 2), consistent with the phenotype of individual *nanT* and *nanX* deletions [21, 24]. TRXC2 expresses all other Sia-utilisation genes (*nanAEKQ, nanCMS, nanY*, and *nagAB* [25, 26]) constitutively due to its parental’s deletions in regulatory genes [19], which means that any specific growth defects will reflect the transporter specificity and/or affinity for the substrate [24, 27, 28].

**Table 1.** Strains and plasmids used in this study.

| NAME | RELEVANT GENOTYPE<sup>a</sup> | SOURCE |
| --- | --- | --- |
| **Strains** |  |  |
| TRXC2 | BW25113($\Delta nanT::FRT, \Delta nanX::F3, \Delta nanR, \Delta nagC$)<sup>b,c</sup> | This work |
| JW5769 | BW25113($\Delta nanY::FRT-Kan^R-FRT$) | [21, 47] |
| **Plasmids** |  |  |
| pWKS30 | Low-copy cloning vector with *lac* promoter and pSC101 *ori* | [48] |
| pES1G | pWKS30 + *nanT* from *Escherichia coli* BW25113 | [49] |
| pES156 | pWKS30 + *nanXY* from *Escherichia coli* BW25113 | [21] |
| pES21 | pWKS30 + *nanX* from *Escherichia coli* BW25113 (ESX1+ESX2) | This work |
| pDRT1 | pWKS30 + *nanG* (B5P37_01885) from *Staphylococcus lutrae* DSM 10244 (ESN601+ESN602) | This work |
| pDRT2 | pWKS30 + *siaX* (B5P37_10005) from *Staphylococcus lutrae* DSM 10244 (ESN603+ESN604) | This work |
| pDRT4 | pWKS30 + *SlnanG*-TEV-His6 | This work |
| pDRT5 | pWKS30 + *SlnanG*[D40A]-TEV-His6 | This work |
| pDRT7 | pWKS30 + *nanX* (STM1132) from *Salmonella typhimurium* LT2 (ESN617+ESN618) | This work |
| pES41 | pWKS30 + *siaT* (STM1128) from *Salmonella typhimurium* LT2 | [24] |
| pSEV21 | pWKS30 + *siaT* (LV622_01385) from *Staphylococcus aureus* RN6390 (ESN358+ESN359) | This work |
| pSEV33 | pWKS30 + *siaX* (FF104_02255) from *Clostridium butyricum* ATCC 19398 (ESN437+ESN438) | This work |
| pKD46 | λRed recombineering functions under *araBAD* promoter control | [50] |
| pCP20 | FLP recombinase expression | [50] |
| pES101 | pUC57kan-F3-*aadA* (Spec<sup>R</sup>)-F3<sup>c</sup> | This work |
| pKD13 | FRT-*nptII* (Kan<sup>R</sup>)-FRT template plasmid | [50] |
| pES134 | Same as pKD13, but with F3-*aadA*-F3<sup>c</sup> | This work<br>Addgene<br>127551 |

b = the *nanR* and *nagC* genes code for the transcriptional repressors of the *nan* and *nag* regulons, respectively [25, 26], hence their deletion results in the constitutive expression of the *nan* and *nag* genes required for Neu5Ac catabolism [25]. See Supplementary Methods for the details of TRXC2 construction.
c = F3 are orthogonal variants of the FRT sites (see Supplementary Methods).

**Figure 2.**
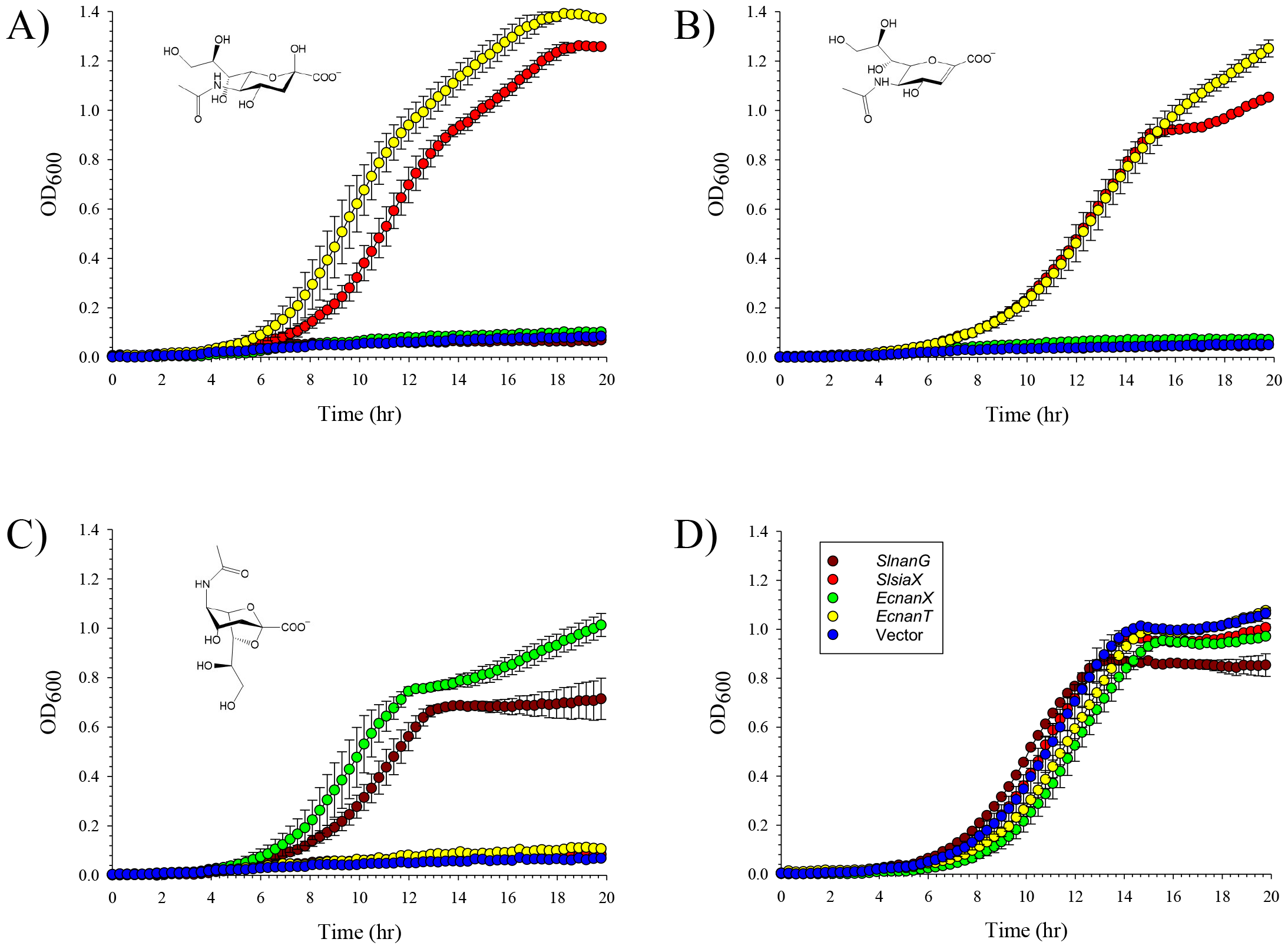
Complementation of the Sia-transport-null strain of *E. coli*, TRXC2, with anhydro-Sia transporter genes. Plasmids carrying different Sia transporter genes under *lac* promoter control were introduced into TRXC2 and the resulting cultures tested for growth in minimal M9 medium with different Sias (structures in the insets) as sole carbon source. Growth experiments were performed in the presence of 1 mg ml^-1^ (≈ 3.5 mM) of either Neu5Ac (A), Neu5Ac2en (B), 2,7-AN (C), or maltose (D), the latter used as a control for Sia-independent growth. Red: *SlsiaX*; brown: *SlnanG*; green: *EcnanX* (positive control for 2,7-AN growth); yellow: *EcnanT* (positive control for Neu5Ac growth); blue: empty vector, pWKS30. Data are the average from triplicate sets ± SD.

To characterise the anhydro-Sia transporters NanG and SiaX, we selected the examples encoded by the genes *B5P37_01885* (*nanG*) and *B5P37_10005* (*siaX*) of *Staphylococcus lutrae* DSM 10244 [29], which were previously computationally identified [17]. Both are linked to NanOx orthologues sharing ≥ 60% identity to the experimentally characterised oxidoreductases, *Rg*NanOx and *Ec*NanY [7, 17, 21]. We tested these transporters alongside *Ec*NanX and *Ec*NanT used as controls (all transporter genes were expressed in TRXC2 from isogenic constructs under IPTG induction; Table 1). Growths of TRXC2 cultures expressing different transporter genes on different Sias as sole carbon source demonstrated that *SlsiaX* mediated uptake of both Neu5Ac and Neu5Ac2en, but not of 2,7-AN (Fig. 2A,B). In contrast, *SlnanG* only supported growth on 2,7-AN, with growth rates and final OD_600_ figures comparable to those conferred by *EcnanX* (Fig. 2C). *EcnanX* also enabled growth on 2,7-AN only, and not on Neu5Ac nor Neu5Ac2en, which instead depended on the presence of *EcnanT* (Fig. 2A,B,C). The growth phenotype of *EcnanX*-expressing TRXC2 was reproduced (Fig. S1) using a different orthologous gene belonging to the same Sia transporter family, the thus-far uncharacterised *STM1132* gene from *Salmonella typhimurium* LT2 [17, 21]. In addition, we confirmed using an isogenic and previously characterised *nanY* mutant of *E. coli* [21], that growth on either 2,7-AN or Neu5Ac2en was dependent on the oxidoreductase EcNanY (Fig. S2).

These results provided experimental evidence for the Sia substrate preference of both uncharacterised transporters, *Sl*NanG and *Sl*SiaX, with *Sl*SiaX being specific for Neu5Ac and Neu5Ac2en, but not for 2,7-AN, and both *Sl*NanG and the control *Ec*NanX transporter being instead specific for 2,7-AN only, as summarised in Table 2.

**Table 2.**
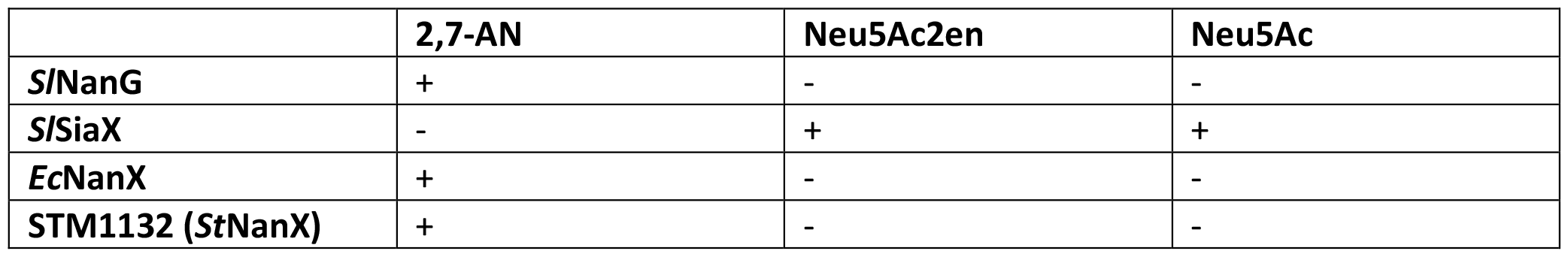
Substrate specificity of the anhydro-Sia transporters investigated in this study.

### A potential 2,7-AN-binding site of *Sl*NanG

We next investigated the structural basis for the substrate preference for 2,7-AN by *Sl*NanG. To identify potential functional residues, we performed a DALI search [30] of the PDB database [31] using an AlphaFold2 [32] model of *Sl*NanG (Fig. 1) as query. One of the best hits was with the crystal structure of *S. typhimurium* MelB, a well-characterised transporter of the GPH (Glycoside-Pentoside-Hexuronide:cation) family, bound with an analogue of the disaccharide substrate melibiose [33] (Fig. S3A). The overlay identified *Sl*NanG D40 (a conserved residue within the NanG/ST8 family; Fig. S3B) as the equivalent of MelB D19 (Fig. S3A), which is a critical substrate-binding site residue forming key H-bonds with melibiose [33, 34]. To investigate the contribution of D40 to NanG function, we generated a *SlnanG* D40A mutant and tested it for complementation of TRXC2 on high (ca. 3.5 mM) and low (ca. 0.7 mM) 2,7-AN concentrations to exacerbate differences. The mutant showed no growth on low 2,7-AN, and poor growth on high 2,7-AN concentration (Fig. 3). The mutation had no effect on protein levels in the *E. coli* cytoplasmic membrane (Fig. S3C). These data are consistent with D40 being a key functional residue of NanG transporters for substrate-binding to 2,7-AN.

**Figure 3.**
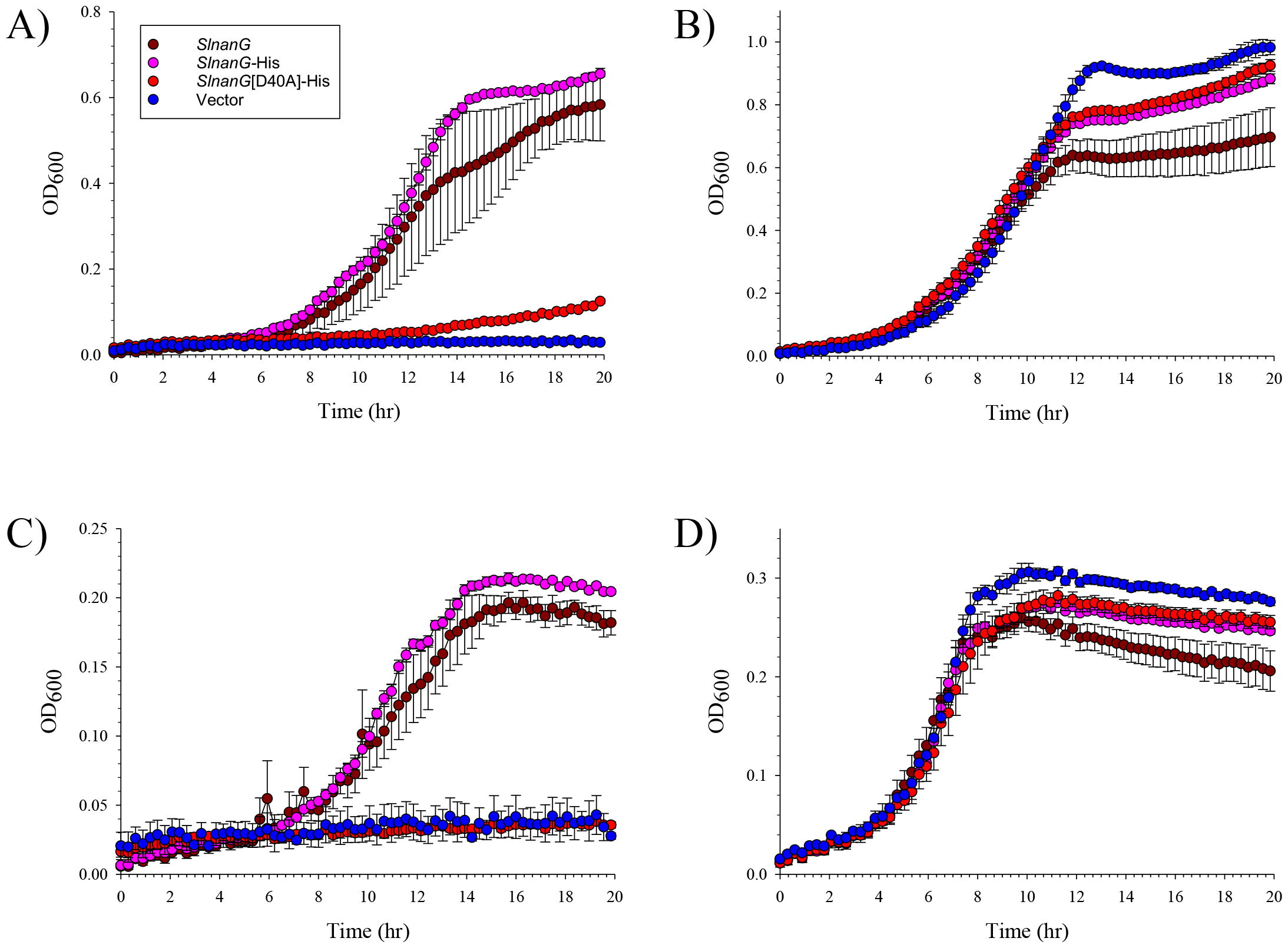
D40 of *Sl*NanG is required for high-affinity uptake of 2,7-AN. Plasmids carrying different alleles of *SlnanG* were introduced into TRXC2 and tested for their ability to complement the growth of this strain on “high” (1 mg ml^-1^ ≈ 3.5 mM; A) and “low” (0.2 mg ml^-1^ ≈ 0.7 mM; C) concentrations of 2,7-AN. B and D: same as in A and C, respectively, but with maltose as control carbon source. Brown: WT *SlnanG*; pink: *SlnanG*-His, coding for a C-terminally His^6^-tagged variant; red: *SlnanG*[D40A]-His; blue: empty vector, pWKS30. Data are the average from triplicate sets ± SD.

### Evidence for distinct substrate specificities in ST5 Sia transporters

SiaT and SiaX transporters of the ST5 family share a conserved Neu5Ac-binding site [17, 35]. SiaT transporters have been extensively characterised for their capacity to transport Neu5Ac [17, 24, 35, 36] and our data showed that *Sl*SiaX can transport both Neu5Ac2en and Neu5Ac substrates. Given the association of NanOx only with some SiaX orthologues [17], we further compared *SlsiaX* with three other ST5 genes unlinked to NanOx, namely, *siaX* (*FF104_02255*) from *Clostridium butyricum* ATCC 19398, and the *siaT* genes from *S. typhimurium* LT2 (*STM1128*) and *Staphylococcus aureus* RN6390 (*LV622_01385*) [37], both coding for characterised Neu5Ac transporters [24, 36]. The variants were tested for growth on high and low concentrations of substrates. All isogenic strains grew comparably well on Neu5Ac and maltose regardless of substrate concentration and transporter expressed (Fig. 4A,B,E,F), but the two *siaX* genes outperformed their *siaT* orthologues in enabling growth on Neu5Ac2en, with growth profiles for Neu5Ac2en resembling those on Neu5Ac even at low concentration of substrate (Fig. 4C,D). These results are consistent with the capacity of SiaX transporters to scavenge both Neu5Ac2en and Neu5Ac, which might underpin their occasional association with NanOx.

**Figure 4.**
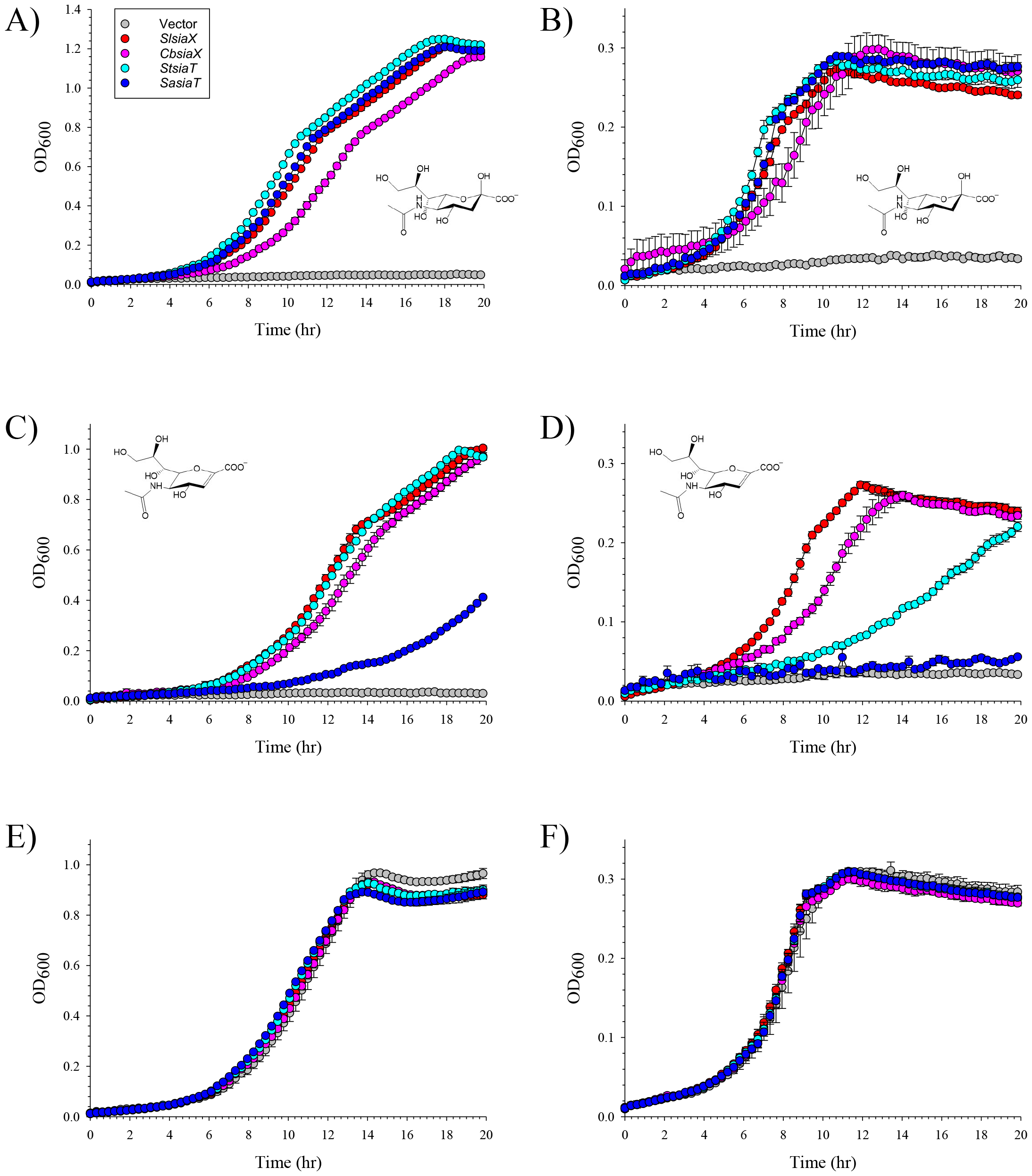
SiaX and SiaT transporters of the ST5 family compared for their ability to take up Sias. Plasmids carrying different ST5 transporter genes were introduced into TRXC2 and tested for their ability to complement the growth of this strain on different substrates at “high” (1 mg ml^-1^ ≈ 3-3.5 mM; A, C, and E) and “low” (0.2 mg ml^-1^ ≈ 0.6-0.7 mM; B, D, and F) concentrations. A and B, Neu5Ac; C and D, Neu5Ac2en; E and F, maltose (control). Red: *SlsiaX*; pink: *CbsiaX*; cyan: *StsiaT*; blue: *SasiaT*; grey: empty vector, pWKS30. Data are the average from triplicate sets ± SD.

## DISCUSSION

Bioinformatic analyses identified four types of Sia transporters involved in anhydro-Sia uptake based on their genetic association with the anhydro-Sia oxidoreductase, NanOx/NanY [17, 21] (Fig. 1). Of these, *Rg*SatXYZ/SAT2 (ST7/ABC) and *Ec*NanX (ST1/MFS) were previously shown to transport 2,7-AN [7, 16, 21]. Here we provided the first experimental evidence for the substrate preference of NanG (ST8/MFS) and SiaX (ST5/SSS) transporters by expressing the *SlnanG* or *SlsiaX* genes in a Sia-transport-null mutant of *E. coli* without interference from the endogenous Sia transporter genes, *EcnanT* and *EcnanX*. We showed that NanG could only transport 2,7-AN, while SiaX was a promiscuous Neu5Ac/Neu5Ac2en transporter.

Previous studies using genetic complementation of single-transporter mutants of *E. coli* showed that *Ec*NanX could transport 2,7-AN only [21], or both 2,7-AN and Neu5Ac2en [16]. Here we showed that, when expressed as the sole Sia transporter in the Sia-transport-null *E. coli* mutant, *Ec*NanX was specific for 2,7-AN, while *Ec*NanT could transport both Neu5Ac and Neu5Ac2en, which is in line with the structural homology of Neu5Ac2en to Neu5Ac but not to 2,7-AN (Fig. 1). In addition, we showed here that a different NanX transporter, the *STM1132* gene product [17, 21], also exclusively transported 2,7-AN. Future biochemical and structural studies of NanX orthologues [17] are warranted to ascertain the 2,7-AN specificity of NanX transporters.

ST5 transporters including some well-characterised SiaT proteins [24, 35, 36, 38] and the SiaX transporter from *C. difficile* [39, 40] function as Neu5Ac transporters. The capacity of SiaX proteins from this family (ST5) to take up Neu5Ac2en efficiently, even scavenging this Sia at growth-limiting concentrations, suggests an additional function in Neu5Ac2en catabolism when associated with cognate NanOx proteins [17]. The sporadic nature of the SiaX-NanOx genetic association [17] suggests however that this might be a partnership of “convenience” which exploits naturally promiscuous transporters to tap into the local pool of Neu5Ac2en released by IT-sialidase action. Indeed, we found that both NanOx-linked *Sl*SiaX and “free” *Cb*SiaX could transport Neu5Ac2en efficiently. Our results from comparing SiaX transporters with the well-known SiaT proteins from *S. typhimurium* LT2 and *S. aureus* RN6390, showed that the latter are poor Neu5Ac2en transporters, and might explain our previous finding that NanOx never forms a partnership with proteins from this group of the ST5 family [17]. Mutagenesis and structural studies are wanted to tease out the details of substrate recognition by different ST5 transporters and to understand the molecular bases of the in-built promiscuity of SiaX transporters.

Overall, our study completed the characterisation of the four types of anhydro-Sia transporters in bacteria. Future studies are needed to investigate the function of the novel transporters, NanG and SiaX, in native organisms to understand the significance of these Sia pathways in host-microbe and microbe-microbe interactions in health and disease.

## METHODS

### Strains and plasmids

Strains and plasmids are listed in Table 1, oligonucleotides are listed in Table S1. Details of strains and plasmids construction are found in Supplementary Methods.

### Growth experiments

For growth experiments using minimal medium, overnight starter cultures of transformed TRXC2 strains were prepared as described previously [24] in M9 medium [41] supplemented with 125 μg ml^-1^ ampicillin (“M9Amp”) and 0.4% v/v glycerol at 37 °C. These were diluted in M9 salts to 10X the required initial OD_600_ (see below) and eventually diluted into experimental M9Amp medium supplemented with 0.1 mM IPTG and carbon source at 0.2-1 mg ml^-1^ (ca 0.7-3.5 mM – see main text for details). Carbon sources were: maltose (Alfa Aesar), replacing glucose [24] for control growth curves, Neu5Ac (Fluorochem), 2,7-AN (made in-house as in [42]), and Neu5Ac2en (Merck). For our initial experiment (Fig. 2) the initial OD_600_ was set at 0.001, but for all later experiments this was increased to 0.01 to shorten the apparent lag phase. Growths were performed in a TECAN Infinite M Nano+ plate reader at 37 °C with shaking every 5 min and sampling every 20 min for 20h. JW5769 transformants were processed in the same manner except that the starter cultures were grown with Neu5Ac 2 mg ml^-1^ instead of glycerol to pre-induce the *nan* regulon [26].

## AUTHORS’ CONTRIBUTION

ES designed the experiments; YW, AB, and ES performed the experiments; TP, FS, and GHT provided resources; ES, NJ, TP, FS, DNB analysed the data; ES and NJ wrote the manuscript with input from TP and DNB; all authors reviewed the final draft of the manuscript.

## ACKNOWLEDGEMENTS

The authors gratefully acknowledge the support of the Biotechnology and Biological Sciences Research Council (BBSRC); this research was partly funded by the BBSRC Institute Strategic Programmes Microbes and Food Safety BB/X011011/1 and Food Microbiome and Health BB/X011054/1. The authors thank Dr Dimitris Latousakis for the ChemDraw structures of sialic acids and Dr Gregor Hagelueken for the AlphaFold2 prediction of the *Rg*SatXYZMsmK transport complex. The authors thank Drs Felicity Alcock and Ben Willson for the kind gift of genomic DNA from, respectively, *Staphylococcus aureus* RN6390 and *Clostridium butyricum* ATCC 19398.

## SUPPLEMENTARY INFORMATION

**Supplementary Methods**

**Supplementary Figures S1-3 legends**

**Supplementary Table S1**

**Supplementary References**

## SUPPLEMENTARY METHODS

### Strains and plasmids construction

Verify (PCRBIO) and Herculase II Fusion (Agilent) were used for high-fidelity PCR, while GoTaq G2 (Promega) for colony PCR. PCRs were treated with DpnI before cloning. Gibson assemblies were performed with either the Quick-Fusion (Biotool) or ClonExpress II (Vazyme) cloning kits. XL1-blue (Agilent) and EC100D*pir*+ (Lucigen/Epicentre) were used as cloning strains.

*E. coli* strain TRXC2 (BW25113Δ*nanT*::FRT, Δ*nanX*::F3,Δ*nanR*,Δ*nagC*,) is a derivative of SEVY3 [1] with the additional in-frame deletion of *nanX*, which was introduced by λRed recombineering [2] followed by marker removal and plasmid curing [3]. The marker cassette, containing the *aadA* (Spectinomycin^R^) gene flanked by F3 sites [4], was amplified from pES134 (construction details below) with KEIO oligos ESN361+ESN353 (yjhB-fwd and -rev from [3]). After genetic modification, all mutant loci of TRXC2 were confirmed by PCR (Table S1) and the new mutation also by sequencing. Strain JW5769 (BW25113Δ*nanY*::FRT-Kan^R^-FRT) from the KEIO collection was used without further modification, as done previously [5].

The pKD13-type template plasmid, pES134, was made by Gibson assembly of PCR products ES184+ES185 for the marker cassette, amplified from the synthetic construct pES101 (GenScript), and ES186+ES187 for the backbone, amplified from pKD13. pES134 reproduces the design of pKD13 [2] and may be used in the same way to introduce unmarked in-frame deletions in the *E. coli* chromosome [3] without illegitimate recombination with pre-engineered FRT scars because of FRT-F3 orthogonality [4]. pES134 is available from Addgene (127551).

Constructs pDRT1, pDRT2, pDRT7, pES21, pSEV21, and pSEV33, all isogenic derivatives of pWKS30 [6] carrying different sialic acid transporter genes under *lac* promoter control, reproduce the design of pES1G, pES41, and pES156, described previously [5, 7, 8], and were made in an equivalent manner by restriction-ligation of PCR products amplified from genomic DNAs (Tables 1 and S1). pDRT4 was made by Gibson assembly of PCR products ESN257+ESN607 for *nanG* (entire coding sequence minus stop codon), amplified from pDRT1, and ESN255+ESN606 for the backbone, amplified from pSEV7 (pSEV7 is a derivative of pWKS30 coding for C-terminal TEV site and His^6^ tag adding sequence - ENLYFQGLEHHHHHH to the last residue of the encoded transporter; Eunice Lee and Emmanuele Severi, unpublished). pDRT5 was made analogously, with the *nanG*-[D40A] insert amplified as two overlapping PCR products (ESN257+ESN612 and ESN613+ESN607). All constructs were verified by sequencing using oligos ESN102 and ESN103 [9].

#### Cell fractionation and western blotting

Transformed TRXC2 strains were precultured as described for the growth experiments, refreshed to an initial OD_600_ of 0.1 in M9Amp supplemented with 0.1 mM IPTG and 2 mg ml^-1^ glucose, and grown to ca 1 OD_600_. Cells were then harvested and subjected to cellular fractionation as described in [9] to separate the membrane fraction, samples of which were then probed with anti-His antibody (Invitrogen) as per the same reference.

### Structural comparison with MelB

Structural models were visualised with CCP4mg [10] or ChimeraX [11]. ChimeraX was also used to superimpose the AlphaFold2 model of *Sl*NanG, generated *via* the Colaboratory online tool [12], with the experimental structure of *Salmonella typhimurium* LT2 MelB-[D59C] bound with the substrate analogue 4-nitrophenyl α-D-galactopyranoside (Protein Data Bank 7L17).

## SUPPLEMENTARY FIGURE LEGENDS

**Figure S1.**
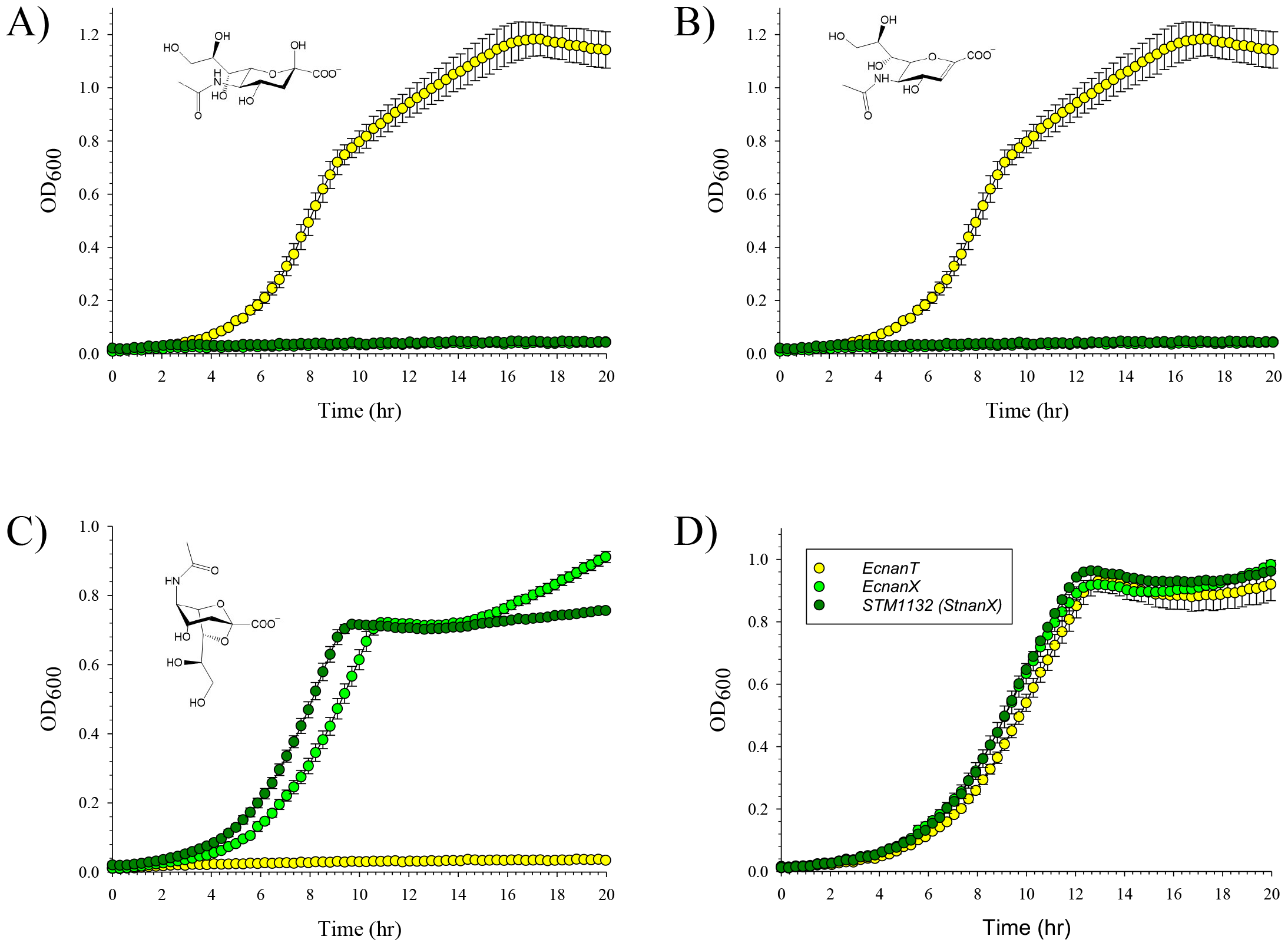
NanX from *Salmonella typhimurium* LT2 is a 2,7-AN-specific transporter. Plasmids carrying different *nanX* genes (Table 1) were introduced into TRXC2 and tested for their ability to complement the growth of this strain on different Sias (all used at 1 mg ml^-1^ ≈ 3.5 mM) as sole carbon source. A: Neu5Ac; B: Neu5Ac2en; C: 2,7-AN; D: maltose. Light green: *EcnanX*; dark green: *STM1132* (*nanX* from *S. typhimurium* LT2); yellow: *EcnanT* (positive control for growth on Neu5Ac, negative control for growth on 2,7-AN). Data are the average from triplicate sets ± SD.

**Figure S2.**
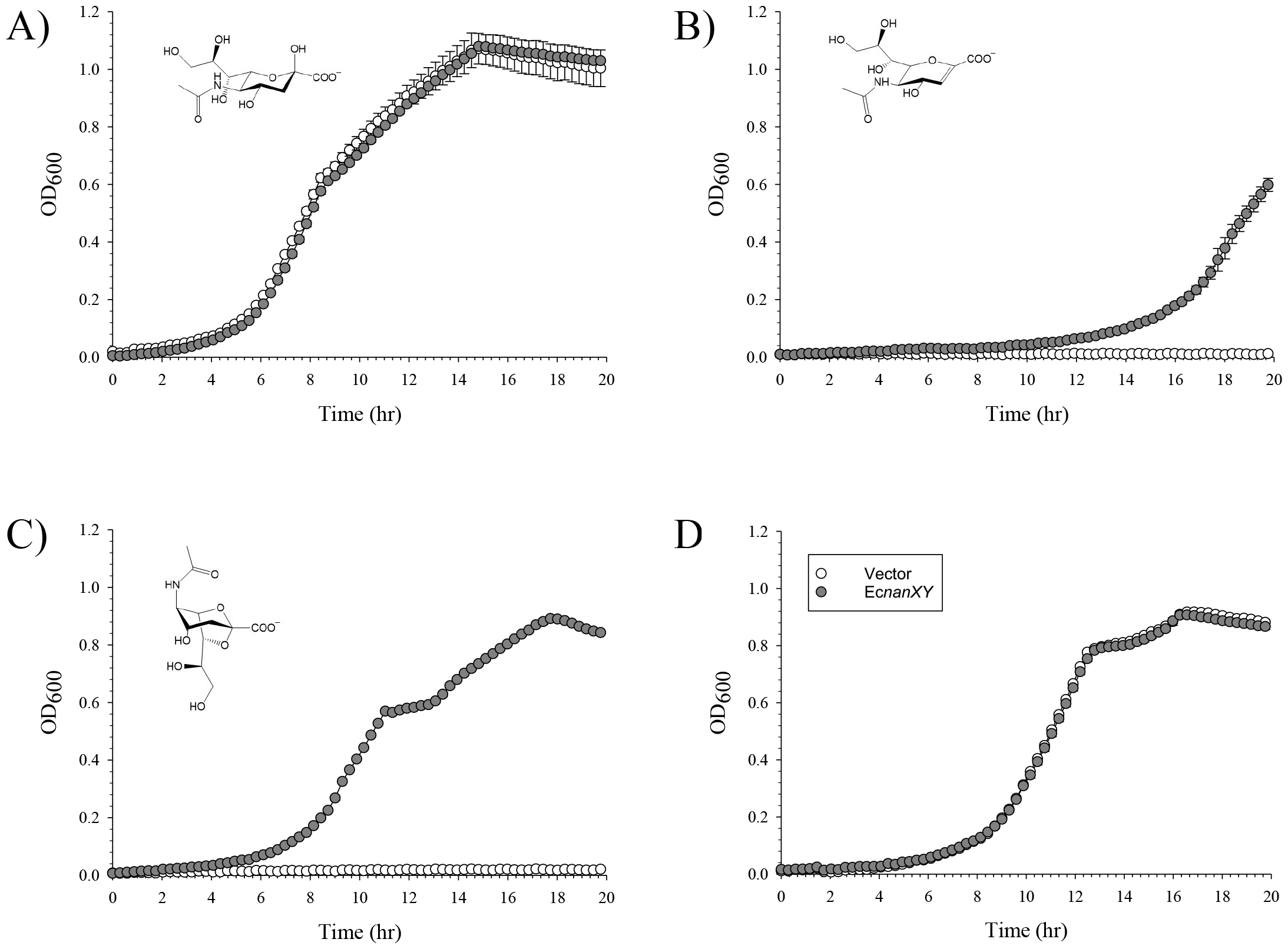
Growth of *E. coli* on anhydro-Sia is dependent on the oxidoreductase NanY. Strain JW5769 (BW25113 Δ*nanY*::FRT-Kan^R^-FRT) was transformed with a plasmid carrying the *EcnanXY* operon (Table 1) and tested for growth on different Sias (all used at 1 mg ml^-1^ ≈ 3.5 mM) as sole carbon source. A: Neu5Ac; B: Neu5Ac2en; C: 2,7-AN; D: maltose. Grey: JW5769 with *EcnanXY*; white: JW5769 carrying empty vector, pWKS30. Data are the average from triplicate sets ± SD.

**Figure S3.**
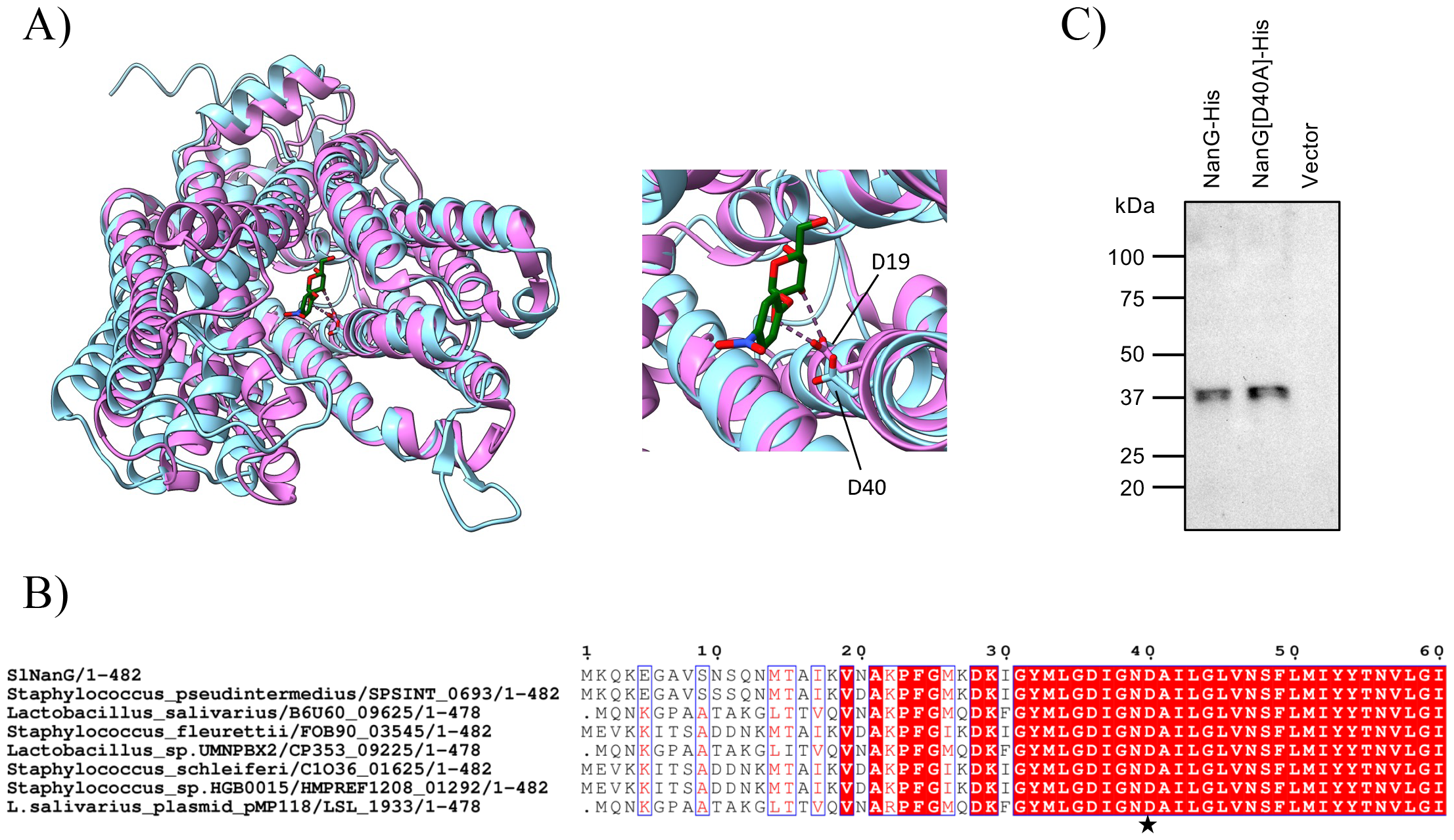
Identification of *Sl*NanG D40 as a functional residue for 2,7-AN uptake. A. Overlay between the AlphaFold2-predicted structure of *Sl*NanG (cyan) and the experimental structure (PDB 7L17) of the melibiose transporter MelB (pink) from *Salmonella typhimurium* (note that AlphaFold2 predicts *Sl*NanG to have 12 transmembrane helices, and not 11 as previously reported [13]). View from the extracytoplasmic side of the transporters into their substrate-binding cavities. Inset: zoom-in on MelB D19 showing the H-bonds with the co-crystallised ligand (the melibiose analogue, 4-nitrophenyl α-D- galactopyranoside) and pinpointing the equivalence of this residue with *Sl*NanG D40; B: Espript [14] alignment (first ∼ 60 residues only) of all NanG transporters of the ST8 family [13], with the asterisk highlighting D40 of *Sl*NanG; C: anti-His-tag immunoblot of membrane fractions from *E. coli* TRXC2 expressing either *Sl*NanG-His^6^ (from pDRT4), the D40A mutant of the same variant (from pDRT5), or no heterologous transporter at all (from pWKS30, “vector”). Notably, *Sl*NanG migrates as a considerably smaller species (37 rather than the predicted 55 kDa), as seen for other membrane proteins [15].

**Table S1.**
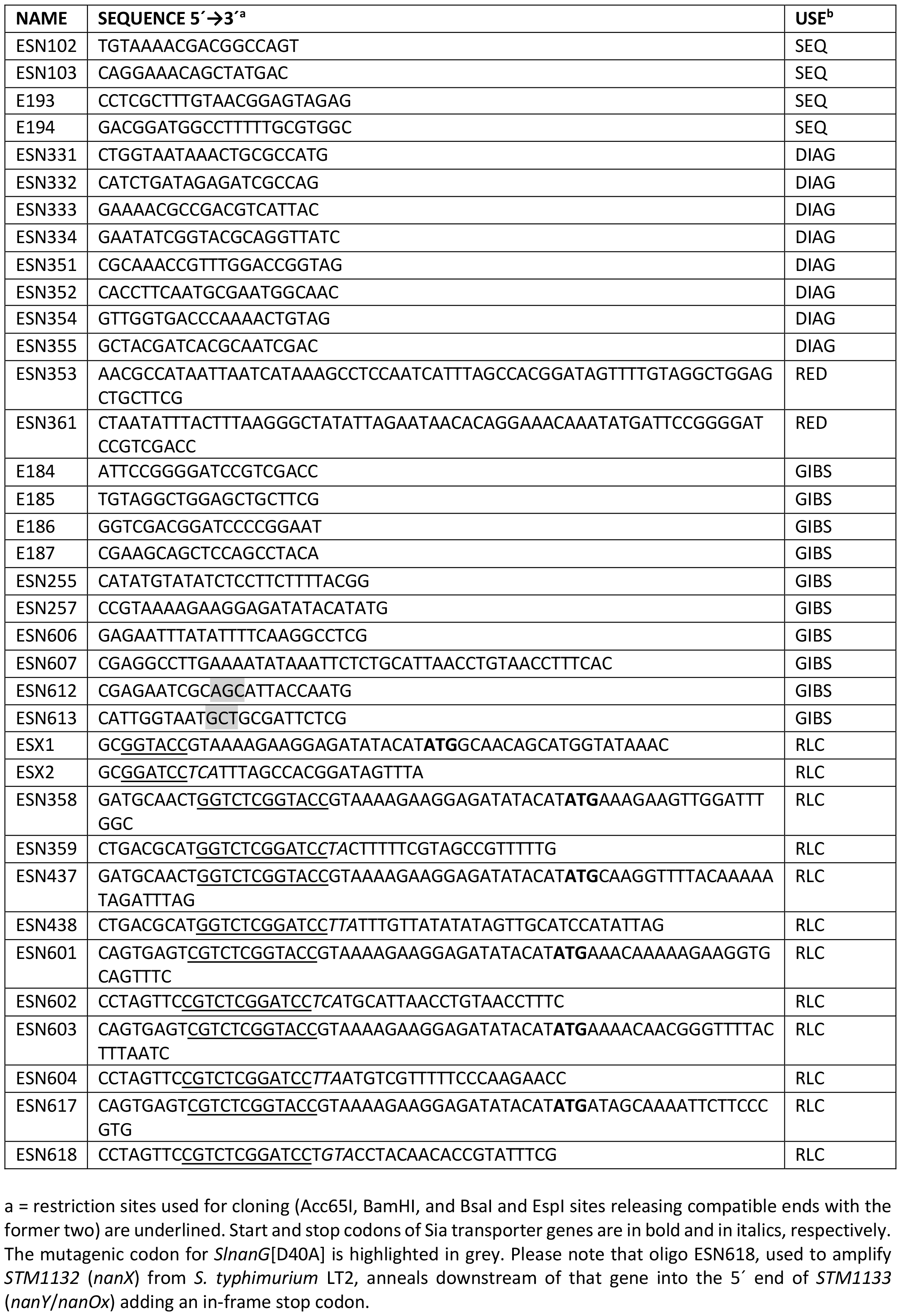
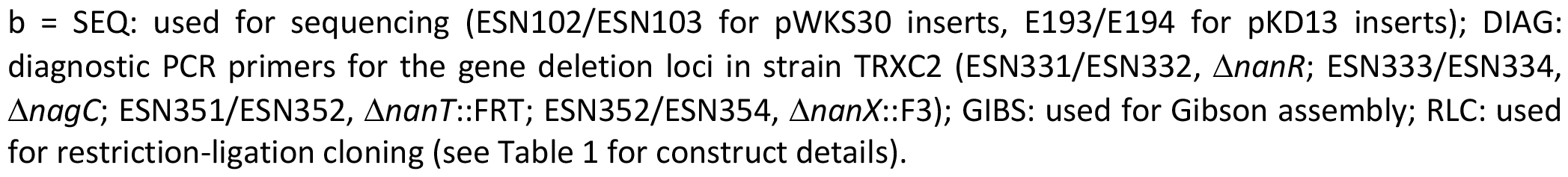

## Notes

### Competing Interest Statement

The authors have declared no competing interest.

## REFERENCES

1. Lewis AL, Chen X, Schnaar RL, Varki A. Sialic Acids and Other Nonulosonic Acids. Epub ahead of print 2022. DOI: 10.1101/glycobiology.4e.15.

2. Varki A. Sialic acids in human health and disease. Trends Mol Med 2008;14:351–360.

3. Angata T, Varki A. Chemical Diversity in the Sialic Acids and Related α-Keto Acids: An Evolutionary Perspective. Chem Rev 2002;102:439–470.

4. Bell A, Severi E, Owen CD, Latousakis D, Juge N. Biochemical and structural basis of sialic acid utilization by gut microbes. J Biol Chem 2023;299:102989.

5. Severi E, Hood DWDWDW, Thomas GHGH. Sialic acid utilization by bacterial pathogens. Microbiology 2007;153:2817–22.

6. Han Z, Thuy-boun PS, Pfeiffer W, Vartabedian VF, Torkamani A, et al. Identification of an N-acetylneuraminic acid-presenting bacteria isolated from a healthy human microbiome. 2020;1–14.

7. Bell A, Brunt J, Crost E, Vaux L, Nepravishta R, et al. Elucidation of a sialic acid metabolism pathway in mucus-foraging Ruminococcus gnavus unravels mechanisms of bacterial adaptation to the gut. Nat Microbiol 2019;4:2393–2404.

8. McDonald ND, Lubin J-B, Chowdhury N, Boyd EF. Host-Derived Sialic Acids Are an Important Nutrient Source Required for Optimal Bacterial Fitness In Vivo. MBio 2016;7:e02237–15.

9. Drula E, Garron ML, Dogan S, Lombard V, Henrissat B, et al. The carbohydrate-active enzyme database: Functions and literature. Nucleic Acids Res 2022;50:D571–D577.

10. Juge N, Tailford L, Owen CD. Sialidases from gut bacteria: a mini-review. Biochem Soc Trans 2016;44:166–75.

11. Xu G, Kiefel MJ, Wilson JC, Andrew PW, Oggioni MR, et al. Three Streptococcus pneumoniae Sialidases: Three Different Products. J Am Chem Soc 2011;133:1718–1721.

12. Tailford LE, Owen CD, Walshaw J, Crost EH, Hardy-Goddard J, et al. Discovery of intramolecular trans-sialidases in human gut microbiota suggests novel mechanisms of mucosal adaptation. Nat Commun 2015;6:7624.

13. Xu G, Potter JA, Russell RJM, Oggioni MR, Andrew PW, et al. Crystal Structure of the NanB Sialidase from Streptococcus pneumoniae. J Mol Biol 2008;384:436–449.

14. Owen CD, Lukacik P, Potter JA, Sleator O, Taylor GL, et al. Streptococcus pneumoniae NanC. J Biol Chem 2015;290:27736–27748.

15. Choi J, Schmukler M, Groisman EA. Degradation of gene silencer is essential for expression of foreign genes and bacterial colonization of the mammalian gut. Proc Natl Acad Sci U S A;119. Epub ahead of print 2022. DOI: 10.1073/pnas.2210239119.

16. Kentache T, Thabault L, Peracchi A, Frédérick R, Bommer GT, et al. The putative Escherichia coli dehydrogenase YjhC metabolises two dehydrated forms of N-acetylneuraminate produced by some sialidases. Biosci Rep 2020;40:1–15.

17. Severi E, Rudden M, Bell A, Palmer T, Juge N, et al. Multiple evolutionary origins reflect the importance of sialic acid transporters in the colonization potential of bacterial pathogens and commensals. Microb Genomics 2021;7:2021.03.01.433349.

18. Thomas GH. Sialic acid acquisition in bacteria–one substrate, many transporters. Biochem Soc Trans 2016;44:760–765.

19. Peter MF, Ruland JA, Depping P, Schneberger N, Severi E, et al. Structural and mechanistic analysis of a tripartite ATP-independent periplasmic TRAP transporter. Nat Commun 2022;13:4471.

20. Davies JS, Currie MJ, North RA, Scalise M, Wright JD, et al. Structure and mechanism of a tripartite ATP-independent periplasmic TRAP transporter. Nat Commun 2023;14:1120.

21. Bell A, Severi E, Lee M, Monaco S, Latousakis D, et al. Uncovering a novel molecular mechanism for scavenging sialic acids in bacteria. J Biol Chem 2020;295:13724–13736.

22. Vimr ER, Troy FA. Identification of an inducible catabolic system for sialic acids (nan) in Escherichia coli. J Bacteriol 1985;164:845–853.

23. Mulligan C, Leech AP, Kelly DJ, Thomas GH. The Membrane Proteins SiaQ and SiaM Form an Essential Stoichiometric Complex in the Sialic Acid Tripartite ATP-independent Periplasmic (TRAP) Transporter SiaPQM (VC1777–1779) from Vibrio cholerae. J Biol Chem 2012;287:3598–3608.

24. Severi E, Hosie AHF, Hawkhead JA, Thomas GH. Characterization of a novel sialic acid transporter of the sodium solute symporter (SSS) family and in vivo comparison with known bacterial sialic acid transporters. FEMS Microbiol Lett 2010;304:47–54.

25. Plumbridge J, Vimr E. Convergent Pathways for Utilization of the Amino Sugars N - Acetylglucosamine, N -Acetylmannosamine, and N -Acetylneuraminic Acid by Escherichia coli. J Bacteriol 1999;181:47–54.

26. Kalivoda KA, Steenbergen SM, Vimr ER. Control of the Escherichia coli sialoregulon by transcriptional repressor NanR. J Bacteriol 2013;195:4689–4701.

27. Fischer M, Hopkins AP, Severi E, Hawkhead J, Bawdon D, et al. Tripartite ATP-independent Periplasmic (TRAP) Transporters Use an Arginine-mediated Selectivity Filter for High Affinity Substrate Binding. J Biol Chem 2015;290:27113–27123.

28. Hopkins AP, Hawkhead JA, Thomas GH. Transport and catabolism of the sialic acids N-glycolylneuraminic acid and 3-keto-3-deoxy-d-glycero-d - galactonononic acid by Escherichia coli K-12. FEMS Microbiol Lett 2013;347:14–22.

29. Veseli IA, Tang C, Pombert J-F. Complete Genome Sequence of Staphylococcus lutrae ATCC 700373, a Potential Pathogen Isolated from Deceased Otters. Genome Announc 2017;5:1–8.

30. Holm L, Laakso LM. Dali server update. Nucleic Acids Res 2016;44:W351–W355.

31. Burley SK, Bhikadiya C, Bi C, Bittrich S, Chao H, et al. RCSB Protein Data Bank (RCSB.org): delivery of experimentally-determined PDB structures alongside one million computed structure models of proteins from artificial intelligence/machine learning. Nucleic Acids Res 2023;51:D488–D508.

32. Jumper J, Evans R, Pritzel A, Green T, Figurnov M, et al. Highly accurate protein structure prediction with AlphaFold. Nature 2021;596:583–589.

33. Guan L, Hariharan P. X-ray crystallography reveals molecular recognition mechanism for sugar binding in a melibiose transporter MelB. Commun Biol 2021;4:1–13.

34. Markham KJ, Tikhonova EB, Scarpa AC, Hariharan P, Katsube S, et al. Complete cysteine-scanning mutagenesis of the Salmonella typhimurium melibiose permease. J Biol Chem 2021;297:101090.

35. Wahlgren WY, Dunevall E, North RA, Paz A, Scalise M, et al. Substrate-bound outward-open structure of a Na+-coupled sialic acid symporter reveals a new Na+ site. Nat Commun 2018;9:1753.

36. North RA, Wahlgren WY, Remus DM, Scalise M, Kessans SA, et al. The Sodium Sialic Acid Symporter From Staphylococcus aureus Has Altered Substrate Specificity. Front Chem 2018;6:1–11.

37. Garrett SR, Mariano G, Palmer T. Genomic analysis of the progenitor strains of Staphylococcus aureus RN6390. Access Microbiol 2022;4:1–11.

38. Bozzola T, Scalise M, Larsson CU, Newton-Vesty MC, Rovegno C, et al. Sialic Acid Derivatives Inhibit SiaT Transporters and Delay Bacterial Growth. ACS Chem Biol 2022;17:1890–1900.

39. Ng KM, Ferreyra JA, Higginbottom SK, Lynch JB, Kashyap PC, et al. Microbiota-liberated host sugars facilitate post-antibiotic expansion of enteric pathogens. Nature 2013;502:96–99.

40. Pereira FC, Wasmund K, Cobankovic I, Jehmlich N, Herbold CW, et al. Rational design of a microbial consortium of mucosal sugar utilizers reduces Clostridiodes difficile colonization. Nat Commun 2020;11:5104.

41. Neidhardt FC, Bloch PL, Smith DF. Culture medium for enterobacteria. J Bacteriol 1974;119:736–747.

42. Monestier M, Latousakis D, Bell A, Tribolo S, Tailford LE, et al. Membrane-enclosed multienzyme (MEME) synthesis of 2,7-anhydro-sialic acid derivatives. Carbohydr Res 2017;451:110–117.

43. Thomas C, Tampé R. Structural and Mechanistic Principles of ABC Transporters. Annu Rev Biochem 2020;89:605–636.

44. Drew D, North RA, Nagarathinam K, Tanabe M. Structures and General Transport Mechanisms by the Major Facilitator Superfamily (MFS). Chem Rev 2021;121:5289–5335.

45. Henriquez T, Wirtz L, Su D, Jung H. Prokaryotic solute/sodium symporters: Versatile functions and mechanisms of a transporter family †. Int J Mol Sci 2021;22:1–21.

46. Marion C, Aten AE, Woodiga SA, King SJ. Identification of an ATPase, MsmK, Which Energizes Multiple Carbohydrate ABC Transporters in Streptococcus pneumoniae. Infect Immun 2011;79:4193–4200.

47. Baba T, Ara T, Hasegawa M, Takai Y, Okumura Y, et al. Construction of Escherichia coli K-12 in-frame, single-gene knockout mutants: the Keio collection. Mol Syst Biol 2006;2:2006.0008.

48. Rong Fu Wang, Kushner SR. Construction of versatile low-copy-number vectors for cloning, sequencing and gene expression in Escherichia coli. Gene 1991;100:195–199.

49. Mulligan C, Geertsma ER, Severi E, Kelly DJ, Poolman B, et al. The substrate-binding protein imposes directionality on an electrochemical sodium gradient-driven TRAP transporter. Proc Natl Acad Sci 2009;106:1778–1783.

50. Datsenko KA, Wanner BL. One-step inactivation of chromosomal genes in Escherichia coli K-12 using PCR products. Proc Natl Acad Sci 2000;97:6640–6645.

## SUPPLEMENTARY REFERENCES

1. Peter MF, Ruland JA, Depping P, Schneberger N, Severi E, et al. Structural and mechanistic analysis of a tripartite ATP-independent periplasmic TRAP transporter. Nat Commun 2022;13:4471.

2. Datsenko KA, Wanner BL. One-step inactivation of chromosomal genes in Escherichia coli K-12 using PCR products. Proc Natl Acad Sci 2000;97:6640–6645.

3. Baba T, Ara T, Hasegawa M, Takai Y, Okumura Y, et al. Construction of Escherichia coli K-12 in-frame, single-gene knockout mutants: the Keio collection. Mol Syst Biol 2006;2:2006.0008.

4. Branda CS, Dymecki SM, Cre- S. Talking about a Revolution: Review The Impact of Site-SpecificRecombinases on Genetic Analyses in Mice. 2004;6:1–22.

5. Bell A, Severi E, Lee M, Monaco S, Latousakis D, et al. Uncovering a novel molecular mechanism for scavenging sialic acids in bacteria. J Biol Chem 2020;295:13724–13736.

6. Rong Fu Wang, Kushner SR. Construction of versatile low-copy-number vectors for cloning, sequencing and gene expression in Escherichia coli. Gene 1991;100:195–199.

7. Mulligan C, Geertsma ER, Severi E, Kelly DJ, Poolman B, et al. The substrate-binding protein imposes directionality on an electrochemical sodium gradient-driven TRAP transporter. Proc Natl Acad Sci 2009;106:1778–1783.

8. Severi E, Hosie AHF, Hawkhead JA, Thomas GH. Characterization of a novel sialic acid transporter of the sodium solute symporter (SSS) family and in vivo comparison with known bacterial sialic acid transporters. FEMS Microbiol Lett 2010;304:47–54.

9. Severi E, Bunoro Batista M, Lannoy A, Stansfeld PJ, Palmer T. Characterization of a TatA/TatB binding site on the TatC component of the Escherichia coli twin arginine translocase. Microbiology 2023;169:1–18.

10. McNicholas S, Potterton E, Wilson KS, Noble MEM. Presenting your structures: The CCP4mg molecular-graphics software. Acta Crystallogr Sect D Biol Crystallogr 2011;67:386–394.

11. Pettersen EF, Goddard TD, Huang CC, Meng EC, Couch GS, et al. UCSF ChimeraX: Structure visualization for researchers, educators, and developers. Protein Sci 2021;30:70–82.

12. Bisong E. Building Machine Learning and Deep Learning Models on Google Cloud Platform. Berkeley, CA: Apress. Epub ahead of print 2019. DOI: 10.1007/978-1-4842-4470-8.

13. Severi E, Rudden M, Bell A, Palmer T, Juge N, et al. Multiple evolutionary origins reflect the importance of sialic acid transporters in the colonization potential of bacterial pathogens and commensals. Microb Genomics 2021;7:2021.03.01.433349.

14. Robert X, Gouet P. Deciphering key features in protein structures with the new ENDscript server. Nucleic Acids Res 2014;42:320–324.

15. Blakey D, Leech A, Thomas GH, Coutts G, Findlay K, et al. Purification of the Escherichia coli ammonium transporter AmtB reveals a trimeric stoichiometry. Biochem J 2002;364:527– 535.

